# Serotonin signals through ciliary 5-HT6 to control insulin secretion

**DOI:** 10.64898/2026.10.30.763155

**Authors:** Shannon E. Townsend, Jeong Hun Jo, Morgan Opp, Katherine Bates, Xinhang Dong, Jing W. Hughes

## Abstract

In the pancreatic islet, serotonin (5-HT) functions basally to modulate glucagon secretion, but its contribution to beta-cell signaling and function remains unclear. Here, we demonstrate that the Gs-coupled serotonin receptor 6 (5-HT6), previously thought to be expressed almost exclusively in the brain, localizes to the primary cilium and plasma membrane of islet cells and augments insulin secretion. Utilizing a transgenic model of beta-cell cilia loss, along with approaches to perturb cilia-specific 5-HT6 function without disrupting the native cilium architecture, we show that 5-HT6 action in beta cells is dependent on the primary cilium. Critically, the plasma membrane pool of 5-HT6 cannot stimulate insulin release when the cilium is absent, representing an instance of whole-cell regulation by a cilia-localized GPCR. These findings establish subcellular localization as a determinant of GPCR function in beta cells and identify ciliary 5-HT6 signaling as a distinct pathway for modulating insulin secretion.

## Introduction

Serotonin (5-hydroxytryptamine, 5-HT) is a monoamine neurotransmitter known for its roles in the central nervous system for regulating mood, appetite, and cognition^1,2^. Outside the brain, serotonin acts as a peripheral signaling molecule, and increasing evidence points to functional roles for serotonin in pancreatic islet biology^3-5^. Serotonin is endogenously produced by islet beta cells, packaged into insulin secretory granules, and co-released with insulin^6^. This co-secretion has been implicated in beta-cell adaptation during pregnancy and puberty, serving as an autocrine signal to enable rapid beta-cell mass expansion and increased insulin secretion, as well as modulating paracrine regulation of glucagon secretion from alpha cells^6-12^. However, the signaling mechanisms by which serotonin regulates normal beta-cell function remain incompletely understood.

Serotonin signals through a family of 15 receptors, most of which are G protein-coupled receptors (GPCRs)^13^. Among these, serotonin receptor 6 (5-HT6) is notable for its predominant expression in the central nervous system and its established roles in learning, memory, and cognition^14^. 5-HT6 is a G_s_-coupled receptor that stimulates cAMP production to activate PKA, though emerging evidence also suggests context-dependent coupling to other G proteins, including G_q_^14,15^. While 5-HT6 has been extensively studied in neurons, its expression and function in peripheral tissues, including pancreatic islets, has remained largely unexplored.

Recent work has revealed that many GPCRs, including incretin and somatostatin receptors^16-18^, localize to the primary cilium – a sensory organelle projecting from the surface of most mammalian cell types that functions as a specialized signaling compartment enriched in GPCRs, ion channels, and second messenger effectors^19-21^. In pancreatic beta cells, we and others have shown that primary cilia play essential roles in glucose homeostasis^22-24^ and that ciliary GPCRs regulate insulin secretion through compartmentalized cAMP and Ca^2+^ signaling^16-18,25^. Whether 5-HT6 localizes to beta-cell cilia and whether ciliary 5-HT6 signaling contributes to the regulation of insulin secretion are unknown.

Here, we demonstrate that 5-HT6 is expressed on the primary cilia of both mouse and human islet cells and, utilizing genetic and pharmacological approaches, we show that 5-HT6 activation stimulates insulin secretion from mouse and human islets. Mechanistically, 5-HT6 signaling elevates PKA activity in the beta-cell cytosol in a cilia-dependent manner, with only modest modulation of cytosolic Ca^2+^, suggesting a distinct signaling mode for ciliary 5-HT6 compared to its well-characterized neuronal counterpart. Crucially, using cilia-targeted loss-of-function approaches, we demonstrate that the ciliary pool of 5-HT6 is specifically required for stimulating insulin secretion, and that the plasma membrane pool cannot compensate. Together, our findings identify 5-HT6 as a ciliary GPCR in pancreatic beta cells and establish that subcellular location dictates its function in the regulation of insulin secretion.

## Methods

### Mice

Ins1-Cre mice from Jackson Laboratories (JAX #026801) were bred with Ift88^fl/fl^ mice (JAX #022409) to generate a beta-cell cilia-deficient mouse line (βCKO) as previously reported^24^. βCKO mice were further crossed with GCaMP6f mice, generously provided by the laboratory of David Piston, producing a double transgenic line bearing both beta-cell cilia knockout and a beta-cell calcium reporter as described previously^16^. Mice were fed a standard rodent diet (PicoLab Mouse Diet 5053, 13.2% calories from fat) and used at 3 to 6 months of age for all experiments. Animals of both sexes were used in studies. Animals were maintained in accordance with Institutional Animal Care and Use Committee regulations at Washington University School of Medicine and Yale University School of Medicine.

### Pancreatic islet isolation and culture

Islets were isolated from adult male and female mice aged three to six months using collagenase digestion and a modified Lacy protocol^24,26^. Following digestion, isolated islets were handpicked into 10-cm plates with 10 mL islet media/plate (RPMI 1640 with 10% fetal bovine serum [FBS], 1% penicillin – streptomycin, and 11 mmol/L glucose) and recovered overnight prior to experiments in a 37°C, 5% CO_2_ incubator. Human donor islets were isolated by and received from the Integrated Islet Distribution Program (IIDP). Upon receipt, human islets were briefly washed in islet media prior to incubation at 37C, 5% CO_2_ in 10-cm plates with 10 mL islet media/plate. Human islets were utilized in studies 24-48h after receipt. Donor information is provided in Supplemental Table 1.

### Immunocytochemistry

Isolated islets were washed with PBS and fixed with 4% paraformaldehyde for 20 minutes, followed by overnight permeabilization and blocking with CAS-Block (Invitrogen) supplemented with 0.3% Triton X-100 (blocking buffer) at 4C. The following day, islets were incubated for 48h in primary antibodies diluted in blocking buffer at 4C. After 48h primary antibody incubation, islets were washed with PBS and subsequently incubated with secondary fluorescent antibodies and dyes (Alexa Fluor, Abberior) diluted in blocking buffer for 24h at 4C. The next day, islets were washed with PBS and incubated for 5 minutes in DAPI (1:10000) to provide nuclear counterstain. Islets were mounted on glass slides with VectaMount AQ Aqueous Mounting Medium (Vector Laboratories) and imaged using Zeiss LSM 880 or Nikon AX/R confocal microscopes. Antibodies and dilutions used were 1:400 ms acetylated tubulin (Proteintech), 1:400 gp ARL13b (Proteintech), 1:400 rb 5-HT6 (Proteintech), 1:400 ms insulin (Proteintech), 1:3 gp insulin (Agilent), 1:1000 ms glucagon (Abcam). Secondary antibodies were ms and gp Alexa Fluor 488/568 and rb Abberior STAR RED dye at 1:500 dilution.

### Pharmacological compounds

All compounds used in studies are commercially available. WAY 208466 dihydrochloride was purchased from Tocris Bioscience (Cat#3904), dissolved in molecular grade water and used at 30 μM concentration in all studies. ST1936 was purchased from MedChemExpress (#HY-103110), dissolved in DMSO and used at 250 nM concentration for live-cell PKA recordings.

### Static hormone secretion assays

Isolated islets (5 per well) were equilibrated in 2.8 mM glucose Krebs-Ringer bicarbonate HEPES (KRBH) buffer (128.8 mM NaCl, 4.8 mM KCl, 1.2 mM KH_2_PO_4_, 1.2 mM MgSO_4_⋅7H_2_O, 2.5 mM CaCl_2_, 20 mM HEPES, 5 mM NaHCO_3_, and 0.1% BSA [pH 7.4]) for 1h at 37C. Following 1h pre-incubation, mouse islets were incubated in 3 mM glucose KRBH (LG condition), or 9 mM glucose KRBH (HG condition) for 1h in the presence or absence of indicated compounds. 2 mM each of L-alanine, L-arginine, and L-glutamine were added to KRBH for LG and HG conditions. Secretion assays for human islets were performed in the same manner except with 12 mM glucose KRBH (HG). At the end of the assay, islets were collected into acid-ethanol solution (2% 12N HCl in 75% ethanol) for islet hormone extraction and stored at -20C. Remaining KRBH containing secreted hormones was collected and stored at -20C until hormone quantification was performed with commercial species-specific insulin and glucagon enzyme-linked immunosorbent assay (ELISA) kits (Crystal Chem, #62100 (mouse), #81518 (mouse), #90095 (human), #81520 (human)). Secreted somatostatin was quantified with a commercially available ELISA kit (Phoenix Pharmaceuticals, #EK-060-03). Hormone secretion data is normalized to hormone content and presented as a percentage of total hormone released.

### Ciliary 5-HT6 delocalization

Tulp3 KD was performed to disrupt ciliary trafficking of 5-HT6. Isolated islets were mildly dissociated with Accutase (StemCell Technologies) for 2 minutes at 37C, washed in PBS, then treated with control adenovirus carrying scramble-shRNA or adenovirus carrying shRNA against mouse *Tulp3* at 1 x 10^8^ viral particles concentration (VectorBuilder) overnight in islet media at 37C, 5% CO2. To assess cAMP dynamics following Tulp3 knockdown, a subset of control and Tulp3 KD islets were simultaneously treated with 1 x 10^8^ viral particles Ad-RIP-R-Flamp1^27^ (VectorBuilder) and incubated for the same duration as islets receiving a single vector. Following a media change the next day, islets were incubated for a further 72h before being utilized in experiments. In brief, glucose-stimulated and 5-HT6-stimulated hormone secretion were assessed as described in the previous section. To visualize cAMP dynamics, scramble and Tulp3 KD islets were plated on recombinant human laminin-521-coated (Gibco) 35 mm glass bottom dishes (Mattek) containing KRBH supplemented with 9 mM glucose, and incubated for at least 30 minutes at 37C, 5% CO2 before imaging. cAMP imaging was performed on a climate-controlled stage maintained at 37°C with a Nikon AX/R point-scanning confocal microscope using the Plan Apochromat λD 10x air objective. Images were acquired at a frame time of 7 seconds/frame. Baseline recordings were performed for approximately 10 min after which islets were treated with 30 uM WAY-208466 and imaged for a further 10 min. At the end of imaging, forskolin was added as a positive control. For analysis, traces were concatenated and normalized to untreated 9 mM glucose condition. Whole-islet ROIs were defined, and mean fluorescence intensity was calculated in Fiji^28^. Data is presented as fluorescence intensity normalized to the 9 mM glucose steady state condition. Secreted hormones and islet insulin content were assayed by species-specific commercial insulin ELISA (Crystal Chem, #62100). Hormone secretion data is normalized to hormone content and presented as a percentage of total hormone released.

To analyze 5-HT6 ciliary localization, following infection with adenovirus and 72h incubation, islets were immunolabeled with acetylated tubulin and 5-HT6 as outlined in Immunocytochemistry. Whole-islet and cilium images were acquired on a Nikon AX/R confocal microscope with the 40x Plan Apochromat λD air objective, with Z-stacks taken through the whole structures. To analyze whole-islet 5-HT6 expression, maximum intensity projections including the maximal islet diameter were created in Fiji^28^, and whole-islet ROIs were defined and analyzed for signal intensity. To analyze ciliary 5-HT6 expression, maximum intensity projections of cilium Z-stacks capturing the whole cilium were created in Fiji^28^, and polygonal ROIs were drawn around each cilium and analyzed for signal intensity of acetylated tubulin and 5-HT6 channels. 5-HT6 ciliary intensity is presented as normalized to acetylated tubulin signal intensity.

### Ciliary PKA disruption

A cilia-targeted PKA inhibitor (PKI) was used to disrupt intraciliary cAMP/PKA signaling. Wild-type mouse islets were transduced with adenoviral constructs carrying Cilia-PKI-4A, cytosol-PKI and Cilia-PKI^29^ (VectorBuilder) at a concentration of 1 x 10^8^ viral particles with the same infection protocol described in the previous section. Islets were subjected to standard hormone secretion assays (described in static hormone secretion assays) in the presence or absence of glucose stimulation and 5-HT6 pharmacologic agonists. Secreted hormones and islet insulin content were assayed by species-specific commercial insulin ELISA (Crystal Chem, #62100). Hormone secretion data is normalized to hormone content and presented as a percentage of total hormone released.

### Live-cell calcium imaging

Wild-type and beta-cell cilia knockout GCaMP6f islets were plated on recombinant human laminin-521-coated (Gibco) 35 mm glass bottom dishes (Mattek) containing KRBH supplemented with 9 mM glucose, and incubated for at least 30 minutes at 37C, 5% CO2 before imaging. Ca^2+^ imaging was performed on a climate-controlled stage maintained at 37°C with a Nikon AX/R point-scanning confocal microscope using the Plan Apochromat λD 10x air objective. Images were acquired at a frame time of 7 seconds/frame. Baseline recordings were performed at 9 mM glucose for approximately 20 min, followed by treatment with 30 uM WAY-208466 and a further 20 min of recording. For analysis, whole-islet regions of interest (ROIs) were defined in Fiji^28^, and traces were concatenated and normalized to baseline 9 mM glucose condition. Oscillation parameters at steady state 9 mM glucose and in the presence of WAY-208466 including oscillation peak value, amplitude, period, plateau fraction and silent phase were quantified using a custom MatLab script (Merrins Lab, https://github.com/hrfoster/Merrins-Lab-Matlab-Scripts)^30^.

### Live-cell PKA and PKC imaging

Wild-type mouse islets and human donor islets were treated with adenoviral constructs Ad-RIP-ExRai-AKAR2^31^ and Ad-RIP-ExRai-CKAR2^32^ (VectorBuilder) at 1 x 10^8^ viral particles concentration in the same manner as previously described studies utilizing adenoviral sensors. On the day of imaging, islets were transferred to recombinant human laminin-521-coated 35 mm glass bottom dishes (Gibco, Mattek) and incubated for at least 30 minutes in 9 mM glucose KRBH before imaging. Islets were imaged in the same manner as described previously. Baseline recordings were performed at 9 mM glucose for approximately 5 min, after which islets were treated with specified compounds and recorded for an additional 10 min followed by treatment with the positive controls forskolin (ExRai-AKAR2) or phorbol 12-myristate 13-acetate (PMA, ExRai-CKAR2). For analysis, traces were concatenated and normalized to untreated 9 mM glucose condition. Whole-islet ROIs were defined, and mean fluorescence intensity was calculated in Fiji^28^. Data is presented as fluorescence intensity normalized to the 9 mM glucose steady state condition.

### Statistical analysis

Data is presented as mean ± SEM unless otherwise reported. Statistical significance was calculated by paired and unpaired student’s t-tests (two groups), and one-way ANOVA or two-way ANOVA (two or more groups) with Tukey’s post-hoc tests where applicable. Sample size and number of replicates for each experiment are included in the figure legends.

## Results

### Endogenous 5-HT6 localizes to primary cilia of mouse and human islets

To determine whether 5-HT6 is expressed in pancreatic islets, we first examined its whole-cell distribution in isolated mouse and human pancreatic islets by immunofluorescence. In both species, 5-HT6 immunoreactivity was detected in both insulin-positive beta cells and glucagon-positive alpha cells, although relatively lower to no expression is observed in alpha cells depending on species (**Fig. 1A, 1B**). Notably, 5-HT6 was detected in a discrete punctate pattern across the cell body and plasma membrane rather than being uniformly distributed, suggesting enrichment in specific subcellular compartments (**Fig. 1A, 1B**).

**Figure 1.**
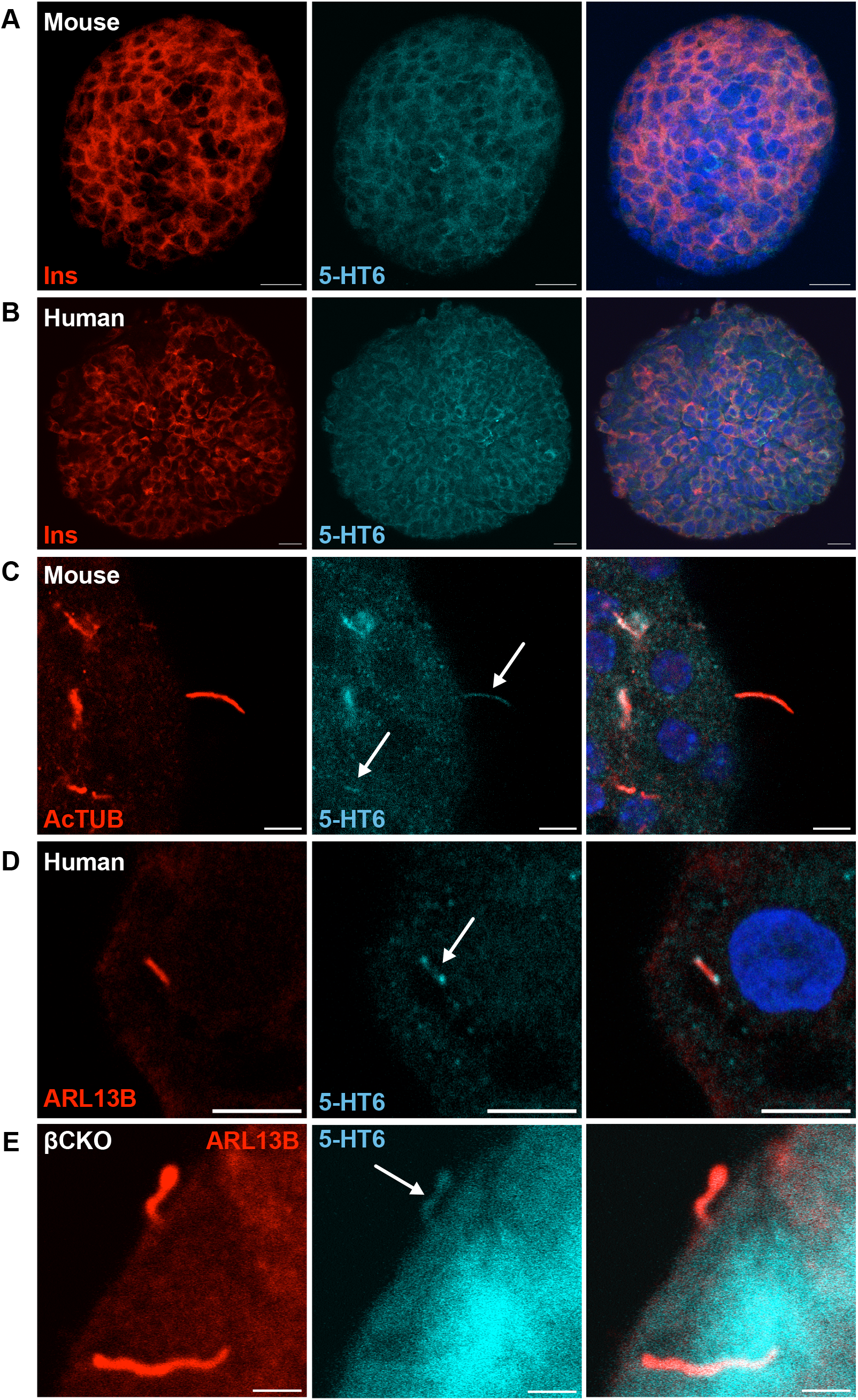
Serotonin receptor 6 (5-HT6) is expressed in mouse and human islets on the plasma membrane and primary cilium. **A**. Wild-type mouse islet immunolabeled for insulin (red) and 5-HT6 (cyan) showing expression of 5-HT6. **B**. Human donor islet immunolabeled for insulin (red) and 5-HT6 (cyan) showing expression of 5-HT6. **C**. Wild-type mouse islet immunolabeled with AcTub (red) to denote cilia and 5-HT6 (cyan), showing 5-HT6 positivity along the ciliary axoneme. **D**. Human donor islet immunolabeled with Arl13b (red) to denote cilia and 5-HT6 (cyan), showing 5-HT6 positivity concentrated at the tip and base of the cilium. **E**. Beta-cell primary cilia-deficient islet immunolabeled for Arl13b (red) to denote cilia and 5-HT6 (cyan), demonstrating non-beta cell primary cilia express 5-HT6. Scale: 20 µm (A and B), 5 µm (C and D), 2 µm (E). White arrows show 5-HT6 ciliary expression.

Given the known enrichment of GPCRs on primary cilia and previously demonstrated 5-HT6 localization to primary cilia in neurons^21,33-35^, we next asked whether 5-HT6 localizes to pancreatic islet cilia. Double immunofluorescence staining for 5-HT6 and the ciliary markers AcTUB and ARL13B revealed colocalization at cilia in both mouse and human islets (**Fig. 1C, 1D**). In mouse islets, 5-HT6 was detected along the length of AcTUB-positive cilia projecting from the cell surface (**Fig. 1C**). Similarly, in human islets, 5-HT6 localized to ARL13B-labeled primary cilia, though the overall cilia length was shorter compared to mouse consistent with previous reports^36,37^. In contrast to mouse islets, 5-HT6 signal was punctate within human cilia, with some seen concentrated near base and tip (**Fig. 1D)**. To assess cell-type specificity, immunolabeling of 5-HT6 in islets from mice with beta-cell-specific knockout of primary cilia (βCKO; IFT88fl/fl; Ins1-Cre)^24^ revealed 5-HT6 localization to non-beta cell cilia, suggesting 5-HT6 is not expressed solely on beta cell cilia (**Fig. 1E**).

Notably, 5-HT6 was also detectable at the plasma membrane in contrast to what has been observed in neurons^33-35,38^, indicating that the receptor is not exclusively ciliary but rather distributed between the two compartments. Further validation of islet 5-HT6 expression has been obtained utilizing 5-HT6-tagged activity reporters which localize strongly to both compartments (**Supp Fig. 1**). Together, these findings demonstrate that 5-HT6 is expressed in mouse and human islets and localizes to the primary cilium, in accordance with its established identity in other tissues as a ciliary protein.

### 5-HT6 activation increases beta-cell PKA and stimulates insulin secretion from mouse and human islets

Having established that 5-HT6 localizes to islet primary cilia, we next asked whether pharmacological activation of 5-HT6 modulates beta-cell signaling dynamics. As 5-HT6 is a G_s_-coupled receptor, agonism should promote cAMP and PKA signaling, which can subsequently influence hormone secretion. To assess how 5-HT6 activation alters PKA dynamics in beta cells, mouse and human islets were infected with ratiometric Ad-RIP-ExRaiAKAR2 (ExRai-AKAR2)^31^, an adenoviral construct carrying a beta-cell-specific cytosolic PKA sensor. Treatment with selective 5-HT6 agonists WAY-208466 (WAY) **(Fig. 2A)** and ST1936 **(Fig. 2B)** increased beta-cell PKA above baseline in both species, demonstrating that 5-HT6 activation stimulates cAMP and downstream PKA activity.

**Figure 2.**
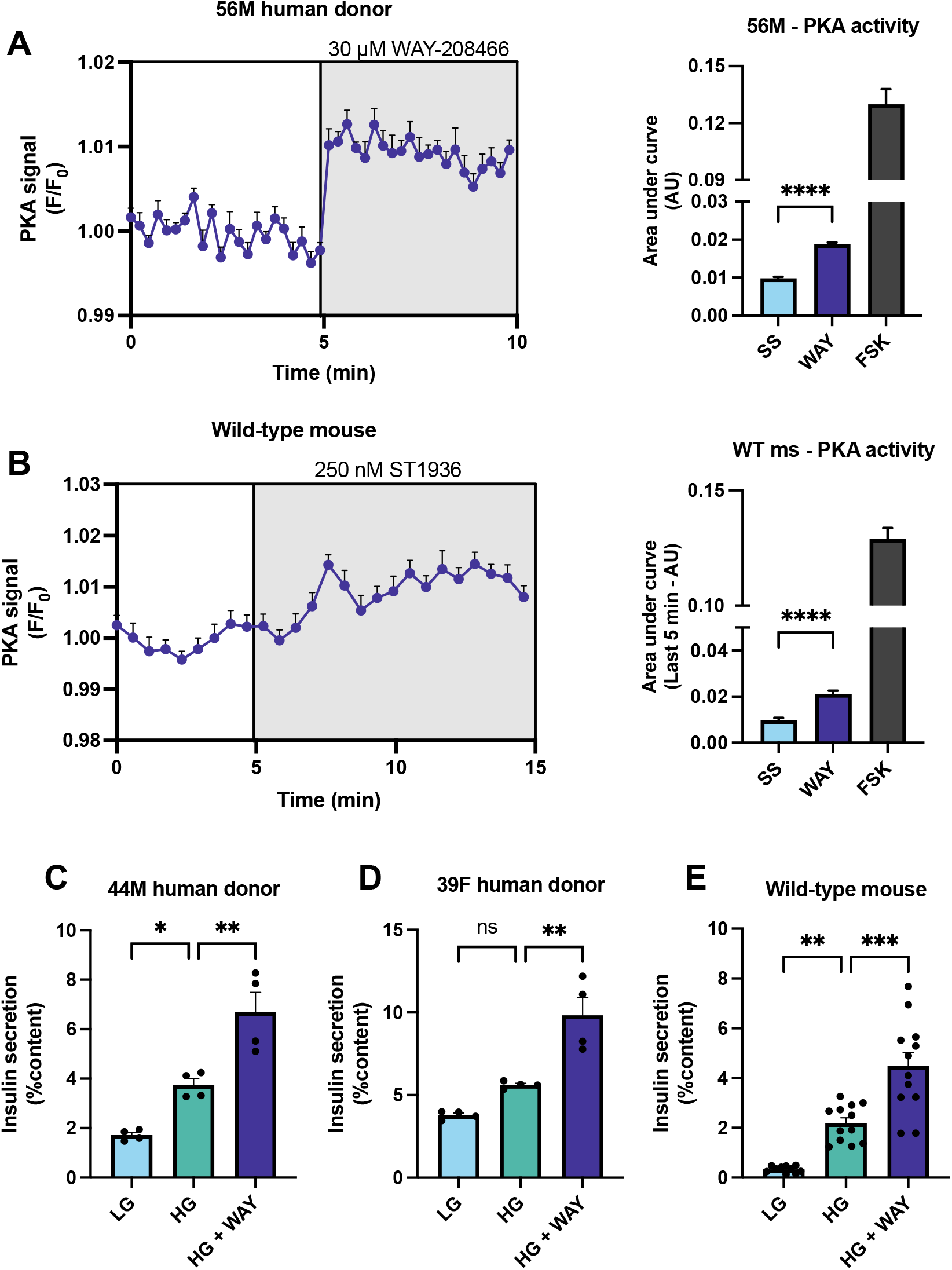
5-HT6 agonism stimulates β-cell PKA activity and potentiates insulin secretion. **A**. Treatment with 5-HT6 agonist WAY-208466 (WAY) in human islets increased beta-cell PKA activity, as measured by ExRaiAKAR2. Area under the curve (AUC) following WAY treatment was significantly greater than the steady state baseline. n = 9 islets. **B**. In wild-type mouse islets, treatment with 5-HT6 agonist ST1936 increased beta-cell PKA activity. AUC following ST1936 treatment was significantly greater than steady state baseline. n = 20 islets. **C**. 5-HT6 agonism (WAY) increases glucose-stimulated insulin secretion in islets from a male human donor. **D**. 5-HT6 agonism (WAY) increases glucose-stimulated insulin secretion in islets from a female human donor. **E**. 5-HT6 agonism (WAY) increases glucose-stimulated insulin secretion in mouse islets, regardless of sex. n = 4 biological replicates. ^*^p<0.05, ^**^p<0.01, ^***^p<0.001, ^****^p<0.0001, ns = not significant.

We next investigated whether stimulation of 5-HT6 can influence hormone secretion. To this end, mouse and human islets were treated with 5-HT6 agonist WAY, and insulin and glucagon secretion were measured under basal (low glucose - 3 mM) and stimulatory (high glucose – 9 or 12 mM) conditions. In human and mouse islets, 5-HT6 agonism significantly augmented insulin release under high glucose conditions (**Fig. 2C-E**) in a sex-independent manner. In line with the finding that 5-HT6 augments insulin secretion, knockdown of 5-HT6 in mouse islets significantly blunted GSIS, demonstrating reliance on the receptor for a robust insulin secretory response to glucose (**Supp Fig. 2**). Furthermore, to assess agonist specificity for 5-HT6, we knocked down the receptor in human islets, resulting in loss of the insulin secretory response to 5-HT6 agonism by WAY (**Supp Fig. 3**). Finally, 5-HT6 agonism significantly increased glucagon and somatostatin secretion at both low and high glucose conditions (**Supp Fig. 4)**, suggesting that 5-HT6 potentiates the secretion of all major islet hormone types.

To determine whether the insulinotropic effect of 5-HT6 requires intact beta-cell cilia, we used βCKO mice harboring beta-cell-specific knockout of primary cilia^24^. In wild-type littermate controls, 5-HT6 agonism robustly increased GSIS as expected (**Fig. 3A, 3B**). However, in male βCKO islets, the potentiating effect of 5-HT6 on insulin secretion was markedly diminished as demonstrated by the stimulation index (**Fig. 3C**). Basal insulin secretion was comparable between control and βCKO islets, whereas the KO islets displayed a select defect in 5-HT6-augmented secretion in high glucose, indicating that cilia loss does not globally impair basal beta-cell function but specifically abrogates 5-HT6-mediated amplification of GSIS. We noted in these functional experiments that cilia loss had a more pronounced effect on 5-HT6-stimulated insulin secretion in male islets compared to females (**Fig 3C, 3D**). Analysis of whole-islet expression of 5-HT6 revealed that female mice express higher levels of the receptor on the plasma membrane compared to male mice (**Fig. 3E**), with whole-islet 5-HT6 signal intensity being significantly higher in females (**Fig. 3F**). Because female islets exhibited higher plasma membrane 5-HT6 expression and a smaller ciliary dependence, we focused subsequent loss-of-function experiments on male islets to isolate the ciliary pool effects. Together, these results establish that 5-HT6 activation potentiates glucose-stimulated insulin secretion from both mouse and human islets, and that this effect requires intact beta-cell cilia.

**Figure 3.**
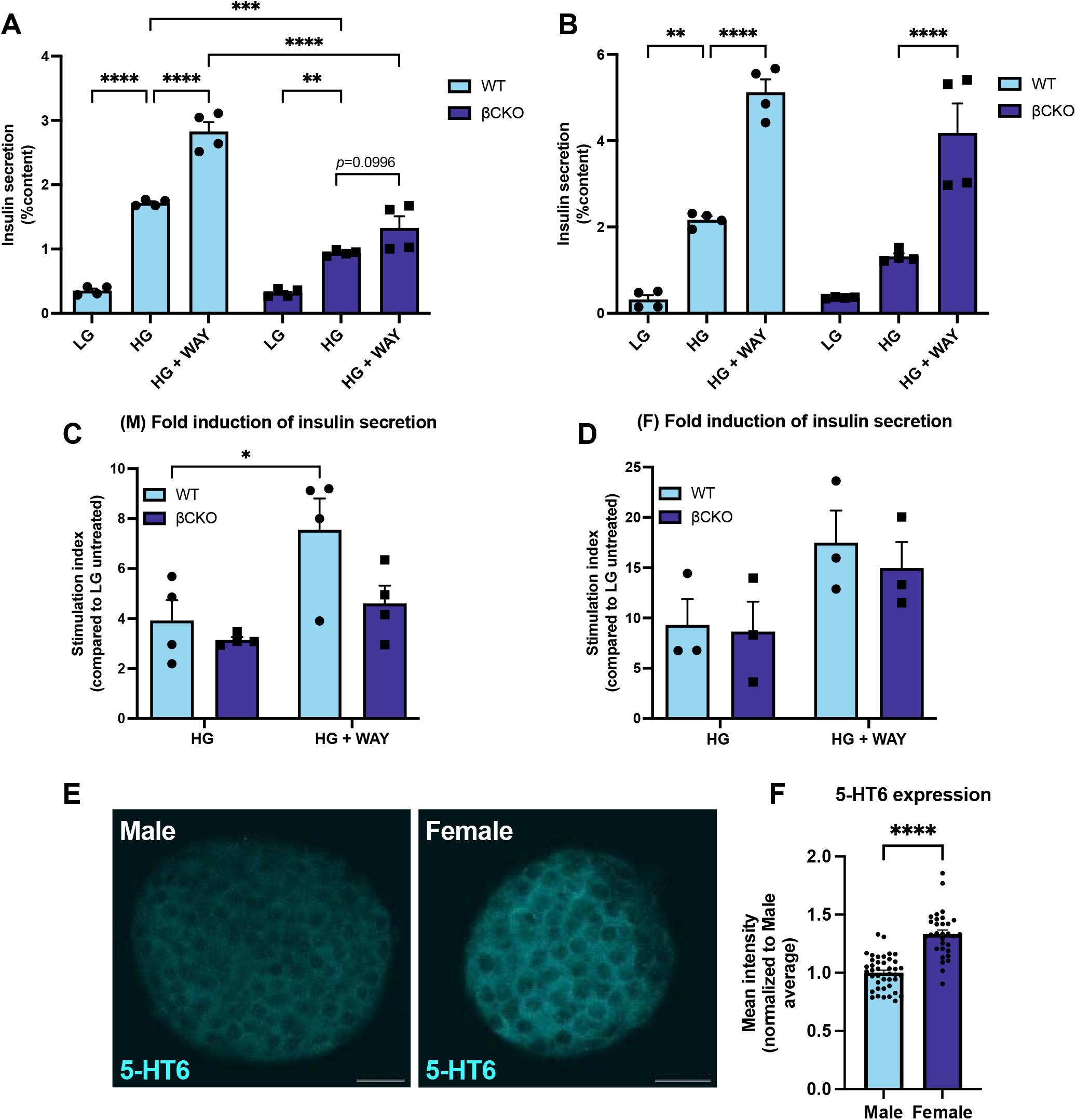
Beta-cell cilia are required for 5-HT6-induced insulin secretion in male mice. **A**. Male wild-type (WT) and beta-cell cilia-deficient (βCKO) islets were treated with 5-HT6 agonist WAY-208466 during a standard hormone secretion assay. Loss of beta-cell cilia significantly blunted 5-HT6-induced insulin secretion. **B**. Female wild-type (WT) and beta-cell cilia-deficient (βCKO) islets were treated with 5-HT6 agonist WAY-208466 during a standard hormone secretion assay. Female βCKO islets retain the ability to secrete insulin in response to 5-HT6 activation. **C**. In male islets, the stimulation index significantly increases following 5-HT6 agonism in only wild-type islets. Stimulation index is calculated from four biological replicates. **D**. In female islets, the stimulation index following 5-HT6 agonism is similar between wild-type and βCKO mice. Stimulation index is calculated from three biological replicates. **E**. Representative image of endogenous staining of a male and female wild-type mouse islet demonstrating sex differences in 5-HT6 whole-islet expression. **F**. Quantification of mean whole-islet intensity of 5-HT6 expression in male and female wild-type islets. Females express significantly more 5-HT6 on the plasma membrane as assessed by immunofluorescence. *n*=41,31 islets for male and female, respectively. ^*\**^*p<*0.05, ^*\*\**^*p*<0.01, ^***^*p*<0.001, ^****^*p*<0.0001, ns = not significant. Scale: 20 µm.

### Ciliary localization of 5-HT6 is dependent on Tulp3 and required for 5-HT6-stimulated signaling dynamics and insulin secretion

While the βCKO model establishes that cilia are necessary, it does not address whether the ciliary pool of 5-HT6 is specifically required, or whether the plasma membrane pool could subserve this function if properly engaged. To address this, we employed two cilia-targeted loss-of-function approaches.

First, we targeted the ciliary adaptor protein Tulp3 known to facilitate 5-HT6 entry into the cilium in other cell types^39^. Furthermore, previous studies have demonstrated that loss of Tulp3 results in partial ciliary delocalization of 5-HT6^39^. To this end, we treated wild-type mouse islets with adenovirus carrying either a scramble-shRNA or an shRNA specific to murine *Tulp3*. Co*-*immunolabeling scramble and Tulp3 KD islets for acetylated tubulin and 5-HT6 revealed significantly decreased ciliary 5-HT6 expression after Tulp3 KD, suggesting that Tulp3 functions to traffic 5-HT6 to the cilium in pancreatic islets (**Fig. 4A, 4B**). Notably, plasma membrane 5-HT6 levels were preserved under Tulp3 knockdown, indicating that the loss of function was specifically attributable to the loss of the ciliary receptor pool rather than global reduction in receptor expression (**Fig. 4C**). Furthermore, acetylated tubulin intensity was similar between the two groups, demonstrating that Tulp3 KD did not significantly disrupt cilium structure (**Fig. 4D**).

**Figure 4.**
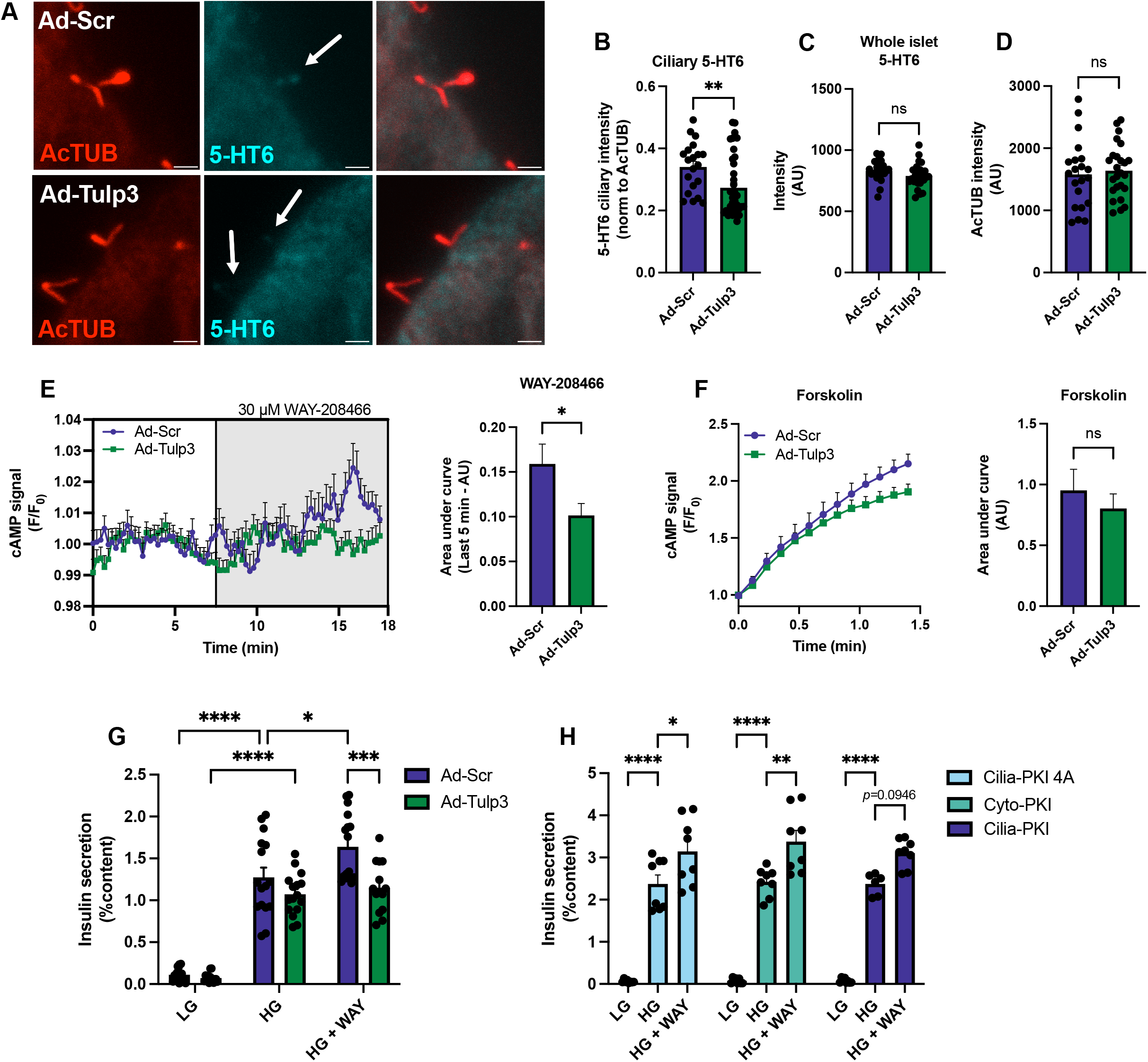
Tulp3 traffics 5-HT6 to islet primary cilia, with ciliary 5-HT6 signaling being necessary for 5-HT6-induced cAMP signaling and insulin secretion. **A**. Representative images of cilia from Ad-Scr and Ad-Tulp3 KD islets. **B**. Quantification of 5-HT6 ciliary intensity normalized to acetylated tubulin intensity. *n*=20 Ad-Scr, 37 Ad-Tulp3 cilia. 5-HT6 ciliary expression is significantly decreased after Tulp3 KD. **C**. Quantification of whole-islet 5-HT6 expression. Tulp3 KD does not alter whole-islet expression of 5-HT6. *n*=23 Ad-Scr, 26 Ad-Tulp3 islets. **D**. Quantification of AcTUB intensity showing no difference between groups. *n*=21 Ad-Scr, 25 Ad-Tulp3 cilia. **E**. Scramble-treated controls (Ad-Scr) or Tulp3 KD (Ad-Tulp3) islets were infected with a beta-cell-specific cAMP sensor and treated with 5-HT6 agonist WAY-208466 (WAY). Tulp3 KD significantly attenuated the cAMP response to WAY, as measured by area under the curve (AUC). *n*=47 Ad-Scr, 42 Ad-Tulp3 islets. **F**. The cAMP response to positive control forskolin is comparable between Ad-Scr and Ad-Tulp3 islets as measured by AUC. **G**. Ad-Scr and Ad-Tulp3 islets were subjected to glucose-stimulated insulin secretion (GSIS) assay with 5-HT6 stimulation. Ad-Tulp3 islets fail to augment insulin secretion in response to 5-HT6 agonism. *n*=4 biological replicates. **H**. Male wild-type islets treated with PKI constructs were subjected to GSIS assay with 5-HT6 stimulation. Loss of ciliary PKA signaling attenuated the insulin secretory response to 5-HT6 agonism. *n*=3 biological replicates. ^*\**^*p<*0.05, ^*\*\**^*p*<0.01, ^***^*p*<0.001, ^****^*p*<0.0001, ns = not significant. Scale: 2 µm.

To assess 5-HT6-stimulated signaling dynamics following Tulp3 KD, scramble (Ad-Scr) and Tulp3 KD (Ad-Tulp3) islets were treated with Ad-RIP-R-Flamp1^27^, a beta-cell-targeted version of a cytosolic sensor measuring cAMP activity. Notably, Ad-Scr and Ad-Tulp3 islets displayed distinct basal cAMP patterns, with Ad-Scr islets displaying small cAMP oscillations which were absent in Ad-Tulp3 islets (**Fig. 4E**). In Ad-Scr islets, treatment with 5-HT6 agonist WAY-208466 (WAY) resulted in an increase in cAMP oscillation amplitude with an eventual rise in cAMP signal above baseline (**Fig. 4E**). In Ad-Tulp3 islets, WAY treatment appeared to cause a few transient cAMP oscillations but did not increase cAMP above baseline (**Fig. 4E**), suggesting that ciliary 5-HT6 is necessary for sustained cAMP activity after receptor agonism. Quantification of the last five minutes of recording demonstrated a significantly higher area under the curve in Ad-Scr islets compared to Ad-Tulp3 islets following WAY treatment (**Fig. 4E**), in line with the diminished cAMP response observed after Tulp3 KD. Treatment with the positive control forskolin increased cAMP signal similarly in each group, demonstrating that knockdown of Tulp3 does not result in a defect in cAMP activation (**Fig. 4F**). Finally, assessment of hormone secretion revealed that Tulp3 KD abolished the insulinotropic effect of 5-HT6 agonism (**Fig. 4G**), suggesting that the ciliary pool of 5-HT6 modulates cAMP/PKA signaling and subsequent insulin secretion downstream of receptor activation.

Second, to directly test whether PKA signaling within the cilium is the relevant downstream effector, we targeted an inhibitory peptide to PKA (PKI) specifically to the ciliary compartment or to the cytosol using GFP-tagged adenoviral constructs. Three constructs were utilized: 1. Cilia-PKI-4A, mutated non-functional PKA inhibitory peptide targeted to the cilium 2. Cilia-PKI, functional PKA inhibitory peptide targeted to the cilium 3. Cyto(solic)-PKI, functional PKA inhibitory peptide targeted to the cytosol^29^. As expected, cilia-PKI-4A islets exhibited 5-HT6-augmented insulin secretion in response to treatment with WAY (**Fig. 4H**). Surprisingly, cyto-PKI islets also responded in the same manner as cilia-PKI-4A islets, suggesting that cytosolic PKA signaling by the receptor is not necessary for 5-HT6-stimulated insulin secretion. In contrast, islets infected with the functional cilia-PKI construct did not display significantly increased insulin secretion in the setting of 5-HT6 agonism (**Fig. 4H**). These findings demonstrate that ciliary PKA signaling downstream of 5-HT6 is required to drive the insulin secretory response, and that the plasma membrane pool of the receptor cannot compensate for its loss.

### Intact beta-cell cilia are required for normal somatic PKA and PKC signaling in response to 5-HT6 activation

To dissect the signaling mechanisms downstream of ciliary 5-HT6 in beta cells, we utilized genetically encoded biosensors to monitor real-time second messenger dynamics in living islets. We first examined PKA activity using a beta-cell-specific version of the cytosol-targeted ratiometric biosensor ExRai-AKAR2^31^. In wild-type mouse islets, stimulation with 5-HT6 agonist WAY-208466 (WAY) induced a sustained increase in PKA activity in the cytosolic compartment as expected (**Fig. 5A**). We next asked whether this PKA response depends on intact beta-cell primary cilia. In βCKO islets, 5-HT6 agonism caused a slight transient increase in PKA activity, with the signal returning to baseline shortly after. Quantification of the area under the curve (AUC) demonstrated a significantly higher PKA response in wild-type compared to βCKO islets **(Fig. 5A)**. Analysis of PKA activity following treatment with the positive control forskolin demonstrated no difference in AUC between the two genotypes, suggesting the blunted response is not due to a global defect in PKA signaling capacity (**Fig. 5B**). Together, these findings indicate that the presence of the primary cilium is required for the sustained somatic PKA response to 5-HT6 agonism.

**Figure 5.**
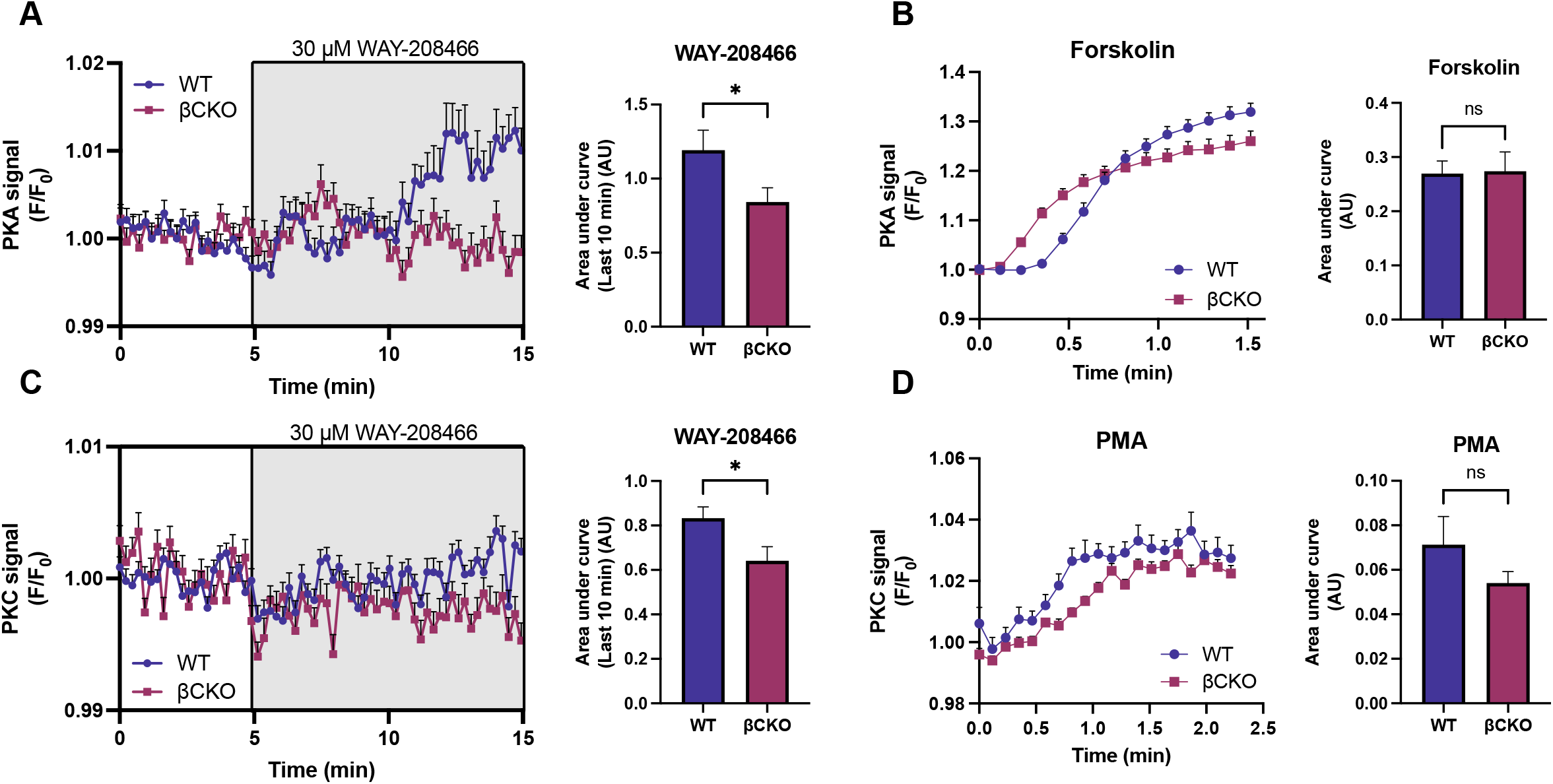
Beta-cell cilia are required for normal somatic PKA and PKC signaling in response to 5-HT6 activation. **A**. Male wild-type (WT) and βCKO islets expressing a beta-cell-specific PKA sensor were treated with 5-HT6 agonist WAY-208466 (WAY). WAY treatment significantly increased PKA signal in wild-type but not βCKO islets as quantified by area under the curve (AUC). *n*=74 WT, 69 βCKO islets. **B**. Male WT and βCKO islets were treated with forskolin as a positive control, with AUC analysis showing no difference in PKA response to forskolin between the two genotypes. **C**. Male WT and βCKO islets expressing a beta-cell-specific PKC sensor were treated with 5-HT6 agonist WAY. AUC analysis demonstrates that βCKO islets have significantly less AUC after WAY treatment compared to WT. *n*=69 WT, 49 βCKO islets. **D**. Male WT and βCKO islets were treated with PMA as a PKC positive control, with AUC analysis showing no difference in PKC response between the two genotypes. ^*\**^*p<*0.05, ns = not significant.

Given that 5-HT6 has been reported to couple to G_q_ in neurons^15^, we assessed whether 5-HT6-G_q_-PKC coupling is present in islets. Wild-type and βCKO islets were infected with adenoviral construct Ad-RIP-ExRai-CKAR2^32^, a beta-cell cytosol-targeted PKC activity reporter. In wild-type islets, agonism of 5-HT6 resulted in a quick transient decrease in PKC signal, followed by a restoration to baseline and gradual rise in signal (**Fig. 5C**). In βCKO islets, WAY treatment resulted in a transient decrease in PKC signal of greater magnitude, with PKC signal remaining below baseline for the remainder of imaging (**Fig. 5C**). Quantification of the area under the curve (AUC) of PKC signal during WAY treatment demonstrated a significantly higher AUC in wild-type compared to βCKO islets. Analysis of the AUC of the PKC response to positive control PMA demonstrated no significant difference between genotypes, suggesting PKC signaling is intact in the absence of beta-cell cilia (**Fig. 5D**). While PKC dynamics were altered in βCKO islets, the lack of a robust PKC response to 5-HT6 agonism in wild-type islets, together with the intact response to PMA, suggests that the Gq-PKC axis is not the primary mechanism by which ciliary 5-HT6 regulates insulin secretion.

We further examined intracellular calcium dynamics using the genetically encoded cytosolic Ca^2+^ indicator GCaMP6f in both wild-type and βCKO mice. In wild-type GCaMP6f islets, 5-HT6 agonism modestly altered calcium oscillation pattern (**Fig. 6A**), causing small but significant decreases in oscillation period and plateau fraction without change to oscillation amplitude, oscillation peak value or silent phase duration (**Fig. 6B**). Surprisingly, despite 5-HT6 activation eliciting only a modest Ca^2+^ response in wild-type mice, this response appeared to be influenced by the presence of beta-cell primary cilia as 5-HT6 stimulation in βCKO GCaMP6f islets altered beta-cell Ca^2+^ dynamics more significantly compared to wild-type (**Fig. 6C**). Similar to wild-type islets, βCKO GCaMP6f islets exhibited significantly decreased oscillation period and plateau fraction, but in contrast had significantly decreased oscillation amplitude, oscillation peak value and silent phase duration (**Fig. 6D**). As βCKO islets still express 5-HT6 on the plasma membrane, this suggests that the presence of the cilium is required for normal whole-cell Ca^2+^ dynamics in response to 5-HT6.

**Figure 6.**
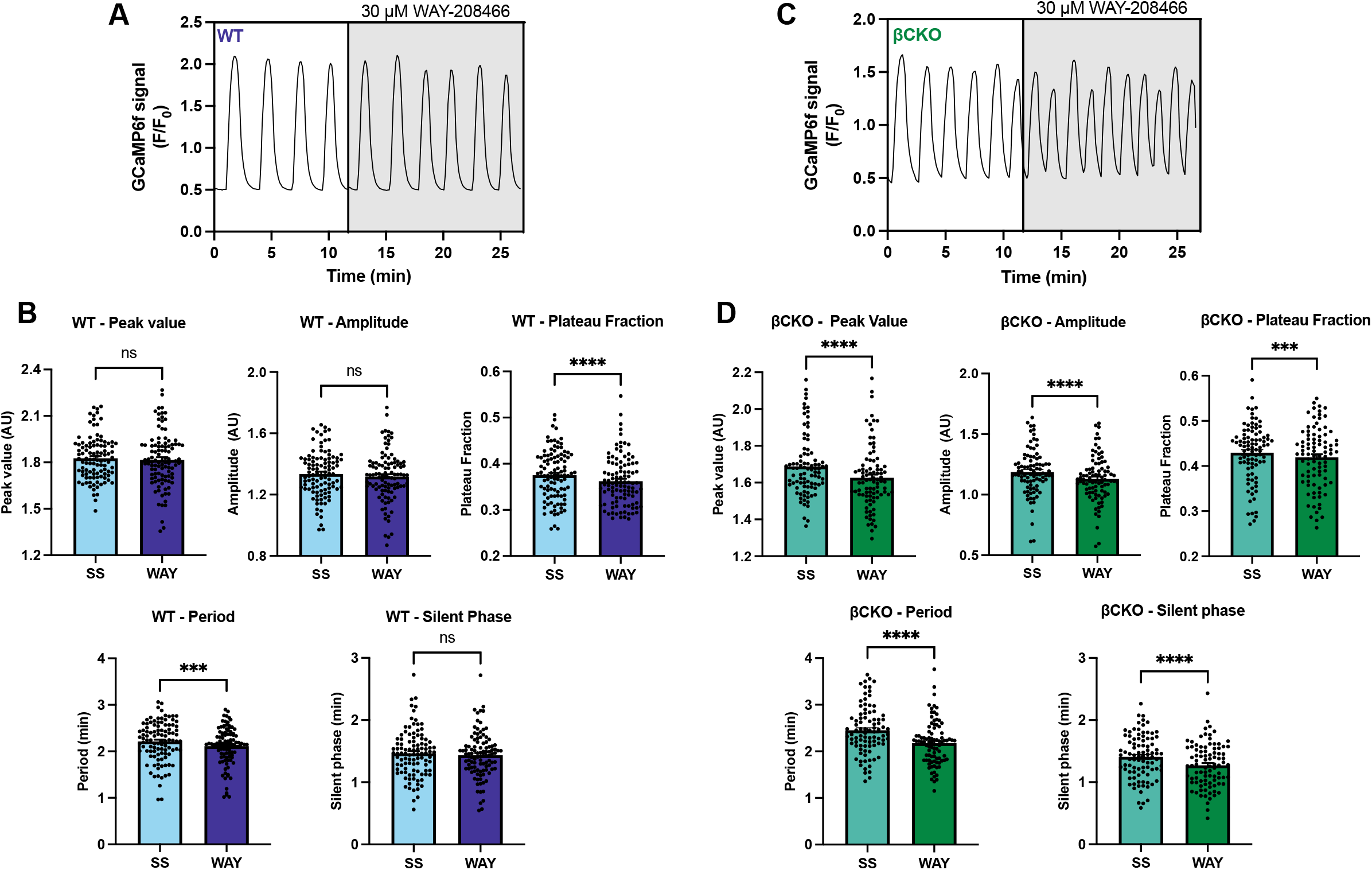
5-HT6 activation modulates whole-cell Ca2+ dynamics in a cilia-dependent manner. A. Representative Ca^2+^ oscillation trace from a wild-type GCaMP6f mouse islet before and after treatment with WAY-208466 (WAY). B. Quantification of Ca^2+^ oscillation parameters from wild-type islets, showing modest but statistically significant reductions in plateau fraction and oscillation period following WAY treatment. C. Representative Ca^2+^ oscillation trace from a βCKO-GCaMP6f mouse islet before and after WAY treatment. D. Quantification of Ca^2+^ oscillation parameters from βCKO-GCaMP6f islets, showing significant reductions across all parameters assessed including oscillation peak value, amplitude, plateau fraction, oscillation period, and silent phase duration. n = 103 WT-GCaMP6f, 89 βCKO-GCaMP6f islets. ^*\**^*p<*0.05, ^***^*p*<0.001, ^****^*p*<0.0001, ns = not significant.

## Discussion

Recently, spotlight has been shone on the primary cilium as more than just a sensory organelle – our group and others have demonstrated that it is a distinct signaling node of the beta cell and crucial for cell function, with dysfunction of beta-cell primary cilia resulting in metabolic defects such as aberrant hormone secretion^22-24^. Furthermore, previous work from our group has highlighted reliance on this organelle for paracrine signaling that is critical for mediating cell-cell crosstalk within the islet^40^. In this report, we present an instance of serotonin signaling through the beta-cell primary cilium modulating beta-cell function. 5-HT6, previously reported to be expressed exclusively in the central nervous system, is expressed in pancreatic islets on the plasma membrane and primary cilium of endocrine cells and mediates hormone secretion from alpha, beta, and delta cells in both mice and humans.

The central conceptual advance of this work is the demonstration that subcellular location dictates the function of 5-HT6 in beta cells. While the receptor is present on both the plasma membrane and the primary cilium, the ciliary pool is specifically required for the insulin secretory response to receptor agonism. The cilia-targeted loss-of-function experiments, including Tulp3 knockdown which preserves membrane receptor levels while depleting the ciliary pool and cilia-PKI which blocks PKA signaling specifically within the cilium, demonstrate that the membrane pool cannot compensate for the loss of ciliary signaling. This finding establishes the cilium as a privileged signaling compartment in which a specific GPCR can drive a physiological output that is not achievable from the plasma membrane. This concept may generalize to other ciliary GPCRs in the islet and beyond.

We have further demonstrated that 5-HT6 action in beta cells is primary cilia-dependent, as loss of beta-cell primary cilia in male mice abolished 5-HT6-induced beta-cell PKA fluxes and insulin secretion. The signaling mechanism downstream of ciliary 5-HT6 is predominantly G_s_-cAMP-PKA, with only modest contributions from Ca^2+^ and little detectable PKC activation, distinguishing it from the neuronal 5-HT6 signaling mode which can engage G_q_-coupled pathways^15^, and suggests that the G-protein coupling profile of 5-HT6 may be context-dependent. Functionally, we show that cilia-localized PKA signaling is required for 5-HT6-stimulated insulin secretion, as direct ciliary PKA inhibition and delocalization of the receptor from the cilium blunts the cAMP and insulin secretory responses in the cell body. In summary, this work establishes 5-HT6 as a cilia-localized GPCR mediating regulation of beta-cell signaling and ultimately function.

A notable observation from our initial characterization was a sex difference in 5-HT6 subcellular distribution in mouse islets. Male islets exhibited lower plasma membrane 5-HT6 expression while retaining robust ciliary 5-HT6 localization, whereas female islets showed higher plasma membrane 5-HT6 expression. We hypothesize that this may confer additional functional reserve for non-ciliary signaling in female islets, supported by the finding that female islets exhibited a smaller reduction in 5-HT6-stimulated insulin secretion after beta-cell primary cilia loss. Given the abundance of the receptor on the plasma membrane and the punctate pattern of labeling, we hypothesize that 5-HT6 likely regulates diverse cellular processes in female islets due to distinct subcellular organelle localization. Serotonin regulation of islet cell function has been historically studied in the context of pregnancy and lactation^7-10,41^, suggesting serotonin biology may be more pertinent to beta-cell function in female mice. The use of only male mice in this study is a limitation to thoroughly understanding 5-HT6 signaling, with future studies aiming to explore sex-dependent signaling by 5-HT6 and define how the receptor regulates islet cell function in a sex-dependent manner.

While the focus of these studies has been characterizing the beta-cell response to 5-HT6 activation, we cannot rule out potential paracrine interactions by glucagon and somatostatin, which are also increased after 5-HT6 activation. Intra-islet glucagon is crucial for potentiation of insulin secretion through binding of GLP-1R receptors on beta cells^42-44^, and our group has reported the presence of a cilia-localized pool of GLP-1R that, when lost, blunts GLP-1R-stimulated cAMP fluxes and insulin secretion^18^. Future studies will assess the potential contribution of glucagon-GLP-1R interactions in the beta-cell response to 5-HT6 activation. Regarding somatostatin, when analyzing PKA signaling, we observed heterogeneous responses to 5-HT6 agonism including both increases and decreases in PKA signal. While the decreases in PKA signal could be attributed to somatostatin secretion in wild-type islets, we have demonstrated that βCKO islets are insensitive to the inhibitory tone of somatostatin due to loss of beta-cell ciliary somatostatin receptor 3 (SSTR3)^16^. While the effects of somatostatin in βCKO islets have been studied in relation to Ca^2+^ signaling, we hypothesize that PKA signaling may be similarly insensitive to somatostatin in the absence of beta-cell primary cilia. This suggests 5-HT6 signaling on the plasma membrane may modulate cellular function in a distinct, non-redundant manner to 5-HT6 on the cilium, which is supported by the Ca^2+^ dynamics assessment where loss of primary cilia significantly altered the Ca^2+^ response to 5-HT6 agonism in βCKO compared to wild-type mice. Future studies utilizing compartment-specific sensors will elucidate the signaling responses elicited after 5-HT6 stimulation on the membrane versus cilium independently, potentially revealing cilia- or membrane-dependent pathways that could be more directly targeted to augment insulin secretion.

To assess the cilia-specific role of 5-HT6, we utilized Tulp3 knockdown to delocalize the receptor from the primary cilium. Prior to this, it was reported that 5-HT6 localization to cilia required Tulp3 in other cell types^39^, and we confirm with this work that Tulp3 traffics 5-HT6 to the cilium in pancreatic islets. 5-HT6 is unique in that it has two cilia localization sequences which each bind Tulp3 or Rabl2 GTPase, both of which are required for complete trafficking of the receptor in neurons and astrocytes^39^. However, based on our functional studies, Tulp3 knockdown was sufficient to abolish the insulin secretory response after 5-HT6 activation, demonstrating potential cell type-specific regulation of receptor localization. While a useful tool, Tulp3 KD is a very blunt tool due to Tulp3 trafficking multiple receptors to cilia including GLP-1R^18,45^. Complementary studies with the cilia-PKI constructs more directly assessed the role of ciliary signaling in 5-HT6-stimulated insulin secretion, demonstrating a blunting in the insulin secretory response when ciliary PKA was inhibited. Surprisingly, while not significant, there was a trend toward increased insulin secretion in the setting of ciliary PKA inhibition, suggesting another potential pathway mediating 5-HT6-augmented insulin secretion. While our studies utilizing the PKC sensor ExRai-CKAR2 measured little change in PKC activity following 5-HT6 agonism, it is possible that this sensor presents a limitation as it has not been extensively studied in islets. Directly assessing contribution of G-protein coupling via pharmacological inhibition of effectors in the G_q_ pathway would provide a clearer understanding of whether 5-HT6-Gq coupling is present in islets.

Finally, the focus of this work has been delineating the function of 5-HT6 in the islet in an *ex vivo* manner. Based on the secretory response elicited from alpha, beta and delta cells upon receptor activation, it is likely that 5-HT6 plays a role in maintaining glucose homeostasis *in vivo*. Previous studies examining loss of 5-HT6 *in vivo* have utilized global inactivation models^46,47^ which may have off-target effects or have conducted virally-delivered Cre recombination to cause acute knockdown of the receptor^35^. Generation of islet cell-type-specific 5-HT6 knockout mouse models will directly allow for assessment of how 5-HT6 modulates islet cell function basally and under pathophysiological conditions. In summary, this work establishes 5-HT6 as a cilia-localized GPCR that regulates beta-cell signaling and function and demonstrates that the ciliary pool of this receptor is functionally required for insulin secretion and cannot be substituted by the plasma membrane pool. These findings expand the known repertoire of ciliary GPCRs in islets and implicate subcellular location as a determinant of GPCR function in beta cells. Future studies leveraging *in vivo* genetic models will be essential to connect the ciliary signaling mechanisms described here to systemic glucose homeostasis.

## Acknowledgements

This work was supported by NIH grants R01DK138974 and R01DK140365 to JWH and the NIH/NIDDK T32 Institutional National Research Service Award to SET (Washington University T32DK007120; Yale University T32DK007058). Human pancreatic islets were provided by the NIDDK-funded Integrated Islet Distribution Program (IIDP) (RRID:SCR_014387) at City of Hope, NIH grant #2UC4DK098085 and the JDRF-funded IIDP Islet Award Initiative.

