## Supplemental Materials for "Serotonin signals through ciliary 5-HT6 to control insulin secretion"

A B

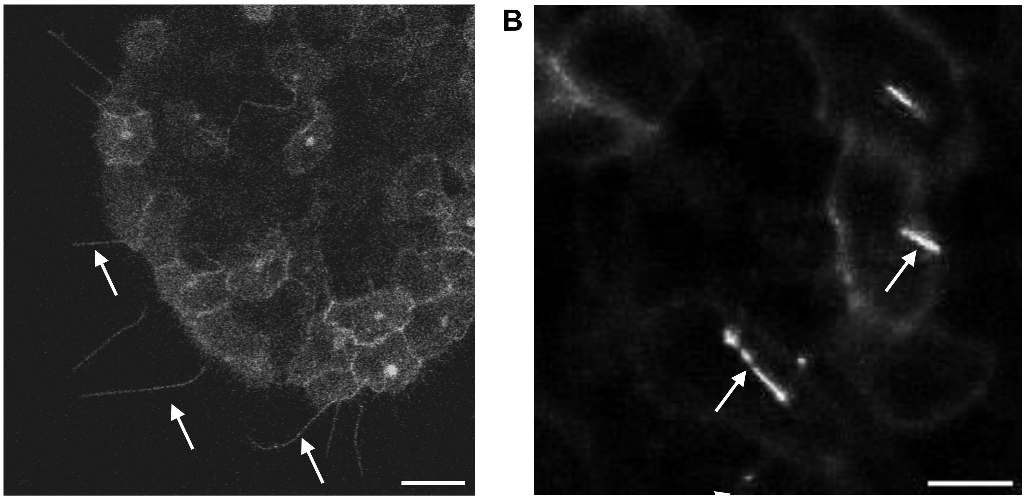

**Supplemental Figure 1. Validation of 5-HT6 expression using an endogenously tagged 5-HT6-PKA reporter. A.** Serotonin receptor 6 (5-HT6) expression in mouse islets detected using Ad-RIP-5HT6-ExRai-AKAR2, an adenoviral construct driving the expression of the PKA sensor ExRai-AKAR2 from the endogenous 5-HT6 locus. Arrows indicate 5-HT6-positive primary cilia. Diffuse 5-HT6 expression is also observed on the beta-cell plasma membrane. Scale: 20 µm. **B.** MIN6 cells were infected with Ad-RIP-5HT6-ExRai-AKAR2. Arrows indicate 5-HT6-positive primary cilia. Scale: 10 µm.

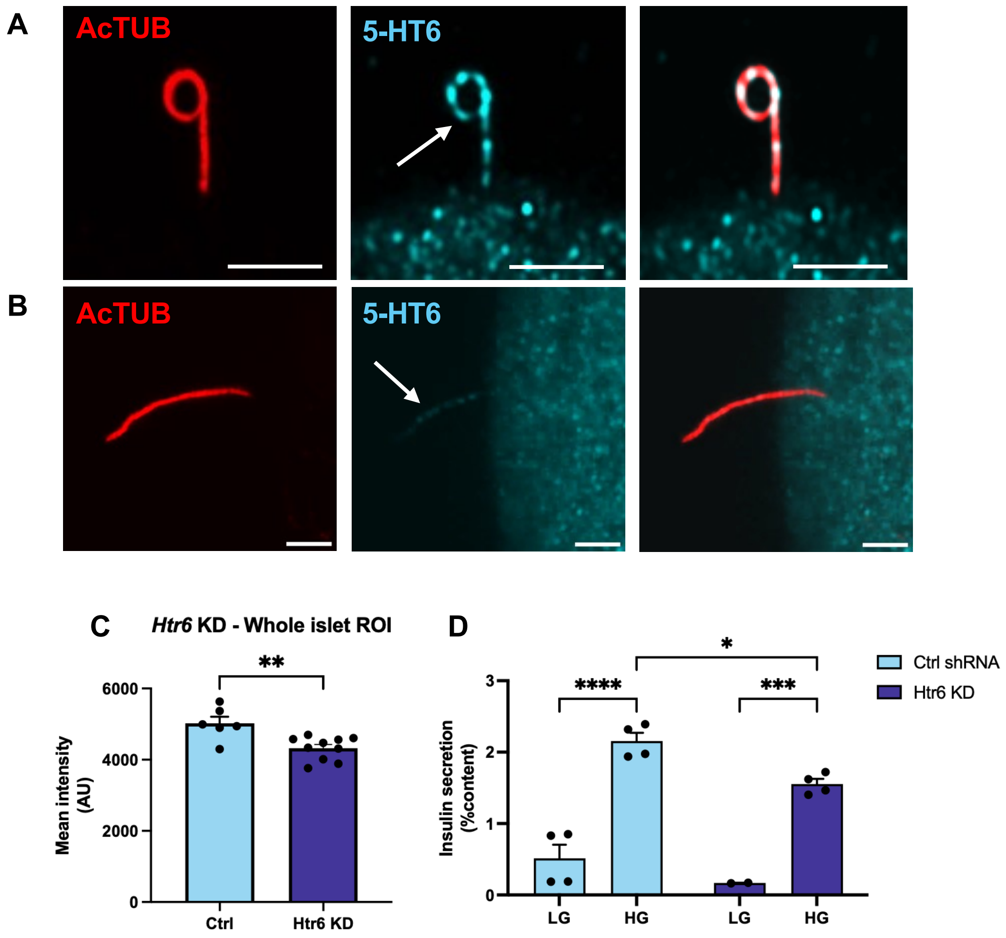


**Supplementary Figure 2. Lentiviral knockdown of 5-HT6 blunts glucose-stimulated insulin secretion (GSIS). A.** Scramble lentivirus-treated mouse islet immunolabeled for AcTUB to mark the cilium (red) and 5-HT6 (cyan). 5-HT6 has punctate expression on the plasma membrane and primary cilium. **B.** *Htr6* knockdown mouse islet immunolabeled for AcTUB to mark the cilium (red) and 5-HT6 (cyan). **C.** Scramble control (Ctrl) and 5-HT6 knockdown (Htr6 KD) whole-islet images were analyzed for mean 5-HT6 labeling intensity. Htr6 KD islets had significantly lower 5-HT6 intensity than Ctrl. **D.** Glucose-stimulated insulin secretion is significantly blunted in Htr6 KD compared to Ctrl. Scale: 2 μm.

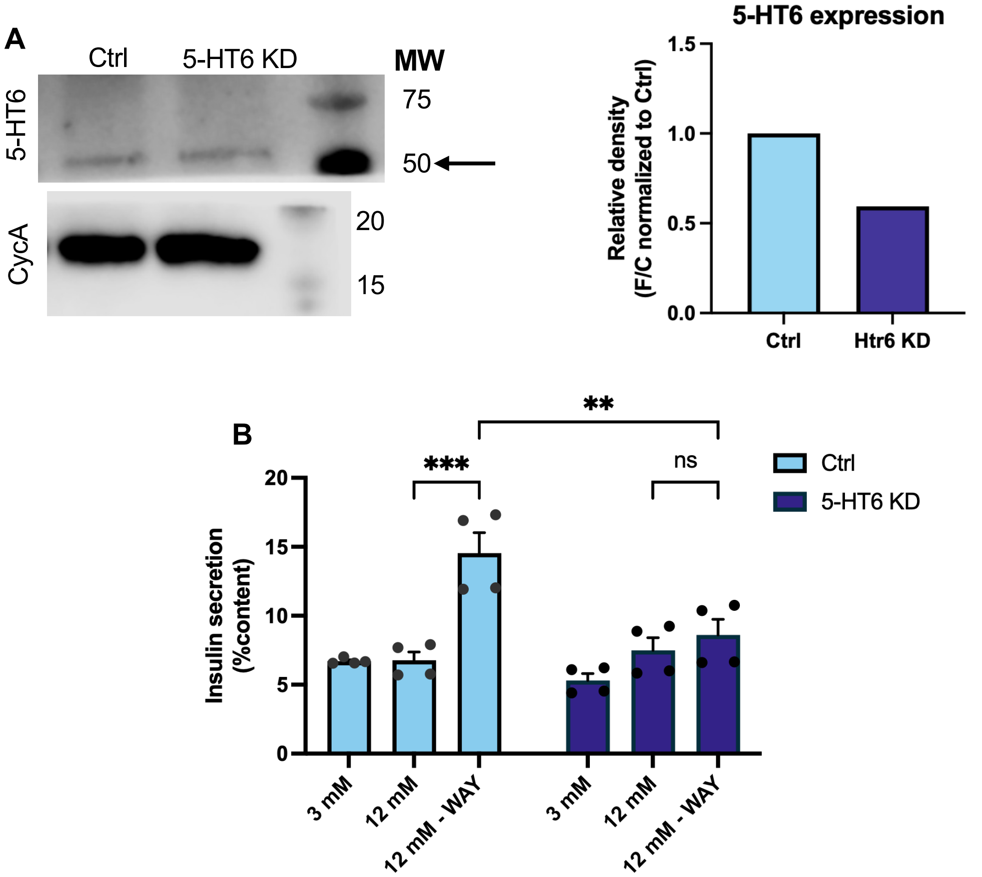


**Supplementary Figure 3. Lentiviral knockdown of 5-HT6 in human islets abolishes the insulin secretory response to 5-HT6 agonism. A.** Western blot for 5-HT6 and cyclophilin A (loading control) expression with densitometry analysis shown. Black arrow denotes 5-HT6 bands, with molecular weight characteristic of human 5-HT6 with post-translational modifications. 5-HT6 protein expression decreased by 40% following receptor knockdown. **B.** Scramble control and 5-HT6 KD islets were subjected to standard hormone secretion assay with 5-HT6 agonism. Knockdown of 5-HT6 abolished 5-HT6-augmented insulin secretion following treatment with agonist WAY-208466. Data are from one human donor. ***p*<0.01, ****p*<0.001, ns = not significant.

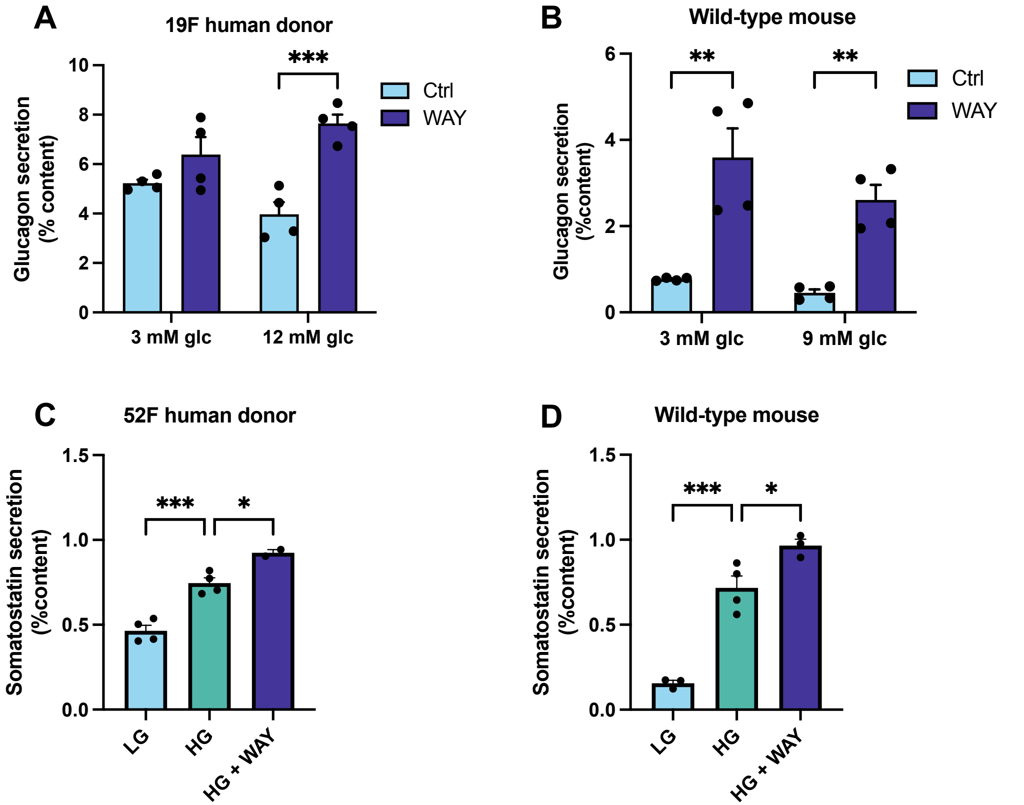


**Supplemental Figure 4. 5-HT6 agonism stimulates glucagon and somatostatin secretion across species. A.** 5-HT6 agonism in human islets increases glucagon secretion at both low (3 mM) and high (12 mM) glucose, with a significant increase under high glucose conditions. **B.** In wild-type mouse islets, 5-HT6 agonism significantly increases glucagon secretion across glucose levels (3 mM and 9 mM).**C.** 5-HT6 stimulation results in somatostatin secretion under high glucose conditions in a human donor. **D.** 5-HT6 agonism induces somatostatin secretion in a wild-type mouse. *$p$<0.05, **$p$<0.01 ***$p$<0.001.

**Supplemental Table 1. Human islet donor characteristics**

| Age | Sex | Race | BMI | HbA1C% | Experiment |
|---|---|---|---|---|---|
| 56 | M | White | 29.1 | 5.4 | PKA |
| 44 | M | Hispanic/Latino | 32.9 | 5.9 | Hormone secretion |
| 39 | F | White | 37.1 | 5.7 | Hormone secretion |
| 19 | F | Black/AA | 22.3 | 6.0 | Hormone secretion |
| 52 | F | White | 30.5 | 5.0 | Hormone secretion, 5-HT6 KD |

**Supplemental Methods.**

**5-HT6 biosensor imaging.**

Isolated wild-type male mouse islets were infected overnight with 1 x $10^8$ viral particles Ad-RIP-5HT6-ExRai-AKAR2, a modified version of the ExRai-AKAR2[31] construct that allows for beta-cell-specific expression of ExRai-AKAR2 where 5-HT6 is endogenously expressed. After a media change the following day and a further 48h of incubation to ensure sensor expression, islets were transferred to a recombinant human laminin-coated (Gibco) glass-bottom dish (Mattek) with 9 mM glucose KRBH imaging buffer (described in main Methods). Live-cell imaging was conducted on a Nikon AX/R point-scanning confocal microscope with the 40x Plan Apochromat λD air objective.

**5-HT6 lentiviral knockdown studies.**

Isolated wild-type male mouse islets were dissociated with Accutase (StemCell Technologies) for 10 min at 37C, then treated with either scramble lentivirus or lentivirus targeted against murine *Htr6* (Sigma; *Htr6* KD sequence: GCGCAACACGTCTAACTTCTT) for 24h in islet media. Following 24h incubation, fresh media was applied and islets were incubated a further 72h in 12-well low-adhesion plates (Thermo Scientific) with media changes every other day. A subset of islets was fixed and immunolabeled for acetylated tubulin and 5-HT6, while the remaining islets were subjected to standard glucose-stimulated hormone secretion assay (described in main Methods). 5-HT6 whole-islet intensity was calculated in Fiji from max intensity projection images including the maximal islet diameters. Secreted insulin and insulin content were measured with species-specific ELISA as described in main Methods.

Human donor islets were dissociated with Accutase (StemCell Technologies) for 20 min at 37C, then treated with either scramble lentivirus or lentivirus targeted against murine *Htr6* (Sigma; *Htr6* KD sequence: CGCCGTCAATTTCTTCAACAT) for 24h in islet media. Following viral infection, islets were handled in the same manner as mouse islets. A subset of islets were lysed in 20 µL of lysis buffer (20 mM Tris-HCl (pH 7.5), 150 mM NaCl, 1 mM $Na_2EDTA$, 1 mM EGTA, 1% Triton, 2.5 mM sodium pyrophosphate, 1 mM beta-glycerophosphate, 1 mM $Na_3VO_4$, 1 µg/ml leupeptin, Cell Signaling) with protease inhibitor (Roche) for 30 min on ice with intermittent vortexing. Lysates were cleared of debris by centrifugation at >10000 rpm for 30 min at 4C. Supernatants were collected and protein content was determined via BCA Assay (ThermoFisher) according to the manufacturer's instructions. Samples were mixed with 4X Laemmli Sample Buffer (Biorad) with 1:10 2-mercaptoethanol (Sigma) and heated at 95C prior to SDS-PAGE analysis. Samples were loaded into 4–15% PAGE mini-PROTEAN TGX gel (Biorad) and electrophoretically separated according to the manufacturer's instructions (100 V for 60 min in Tris/Glycine/SDS Running Buffer, Biorad). Separated proteins were transferred to nitrocellulose membrane 0.2 µm (Biorad) for 90 min at 100 V in transfer buffer (Tris/Glycine 10% Methanol) at 4C. Membrane was exposed to blocking buffer (TBS-tween with 1.5% BSA) overnight at 4C and then primary antibodies (1:1000 cyclophilin A, Proteintech; 1:500 5-HT6, Bioss) overnight at 4C. Membranes were washed with TBS-T and then exposed to secondary antibody (1:2000 anti-rabbit Igg HRP, Abcam) for 1 hour at room temperature followed by additional washes with TBS-T. Chemiluminescent substrate (SuperSignal West Pico PLUS, ThermoFisher) was applied per manufacturer instruction and membranes were imaged using LI-COR Odyssey Fc imaging system. Densitometry analysis was conducted in LICORbio Image Studio software. Remaining islets were subjected to standard glucose-stimulated hormone secretion assay (described in main Methods) in the presence or absence of 30 uM 5-HT6 agonist WAY-208466. Secreted insulin and insulin content were measured with species-specific commercial ELISA (Crystal Chem, #62100 (mouse), #90095 (human)).

**Static islet imaging.**

Mouse islets following lentiviral knockdown were fixed and immunolabeled for AcTUB and 5-HT6 as outlined in the main Methods (Immunocytochemistry). Confocal imaging was performed using a Nikon AX/R point-scanning confocal microscope equipped with Nikon spatial array confocal (NSPARC) detector. Images were acquired using Plan Apochromat 60x and 100x silicone oil immersion objectives (NA 1.35). All imaging used the Galvano scanner with the following parameters: 2 μs dwell time, 2x scan averaging, and 2048 x 16-pixel resolution. Laser power and z-stack settings were optimized for each channel to maintain consistent signal intensity throughout the imaging volume while using approximately 30% of the detector's dynamic range.
